# Regression-based Modeling of Spearman’s Rho for Longitudinal Metabolomics and Mental Wellness in Breast Cancer Patients

**DOI:** 10.64898/2026.04.13.718341

**Authors:** Y. Chen, T.T. Gui, Z. Huang, N.E. Quach, S. Tu, J. Liu, T.J. Garrett, A.R. Starkweather, D.E. Lyon, B.E. Shepherd, X.M. Tu, T. Lin

## Abstract

Chemotherapy in breast cancer (BC) can substantially affect mental wellness. Advances in metabolomics enable comprehensive profiling of metabolic changes over time during and after treatment, offering insights into biological mechanisms linking chemotherapy to mental health outcomes. To study the association between metabolite profiles and mental wellness, correlation-based analyses are particularly useful. Spearman’s rho is a widely used correlation measure and popular alternative to Pearson’s correlation, since it also applies to non-linear association between variables. However, existing methods are not designed for longitudinal data and do not allow for covariate adjustments. In this paper, we propose a novel regression-based framework grounded in a class of semiparametric models, the functional response models, to extend this popular correlation measure to longitudinal settings with missing data under the missing at random assumption. This framework facilitates inferences about temporal changes in correlations over time and association of explanatory variables for such changes. We use simulation studies to evaluate performance of the approach with moderate sample sizes. We apply the approach to a one-year longitudinal substudy of the EPIGEN study to examine the longitudinal association between metabolite profiles and mental wellness in BC patients undergoing chemotherapy. The identified metabolites may serve as candidates for future in-depth bioinformatics analyses and translational investigations.

## 1. Introduction

Breast cancer (BC) is the most common cancer diagnosed in women worldwide, and its incidence has continued to increase in recent years (Siegel *et al*., 2025; World Health Organization, 2025). BC treatment often involves chemotherapy that can substantially impact both physical and mental well-being (Carreira *et al*., 2018; Reece et al., 2013). Recent innovations in metabolomics, the global profiling of small-molecule metabolites such as amino acids, organic acids, and acylcarnitines, have enabled comprehensive assessment of how metabolism changes over time during and after cancer treatment (Jacob *et al*., 2019). Metabolomics and metabolic pathways provide an important framework for understanding the impact of chemotherapeutic agents on cellular responses that in turn can influence wellness (Schmidt *et al*., 2021; Kar et al., 2024). Therefore, identifying longitudinal associations between metabolite profiles and mental health outcomes in patients with BC receiving chemotherapy could provide insight on potential therapeutics targets to optimize mental wellness. When studying such associations, changes in metabolite levels may either influence or result from changes in mental health. Moreover, metabolomics features are often highly correlated and can introduce multicollinearity, overfitting and bias in regression models with multiple features. Therefore, correlation-based analyses are more suitable in the initial stage of biomarker discovery to identify a focused set of candidate metabolites, consistent with approaches proposed in the feature-screening literature (Fan and Lv, 2008; Li, Zhong and Zhu, 2012).

Correlation is widely used to assess strengths of association between variables. The Pearson correlation is arguably the most popular measure of correlation in research and practice for continuous outcomes. One major limitation of this popular measure is that it is a useful index of correlation only when measuring linear relationships between the variables. If the relationship is non-linear, the Pearson correlation generally does not provide a good indication of association between the variables and can lead to misinformed conclusions. Spearman’s rho is a popular alternative for assessing correlation. Unlike its Pearson counterpart, Spearman’s rho does not require linear relationship and provides a measure of correlation for a broader class of monotonic relationships.

This work is motivated by a longitudinal substudy of the EPIGEN, which examined associations between multiple biological and psychological exposures and cancer treatment over time (see Section 2 for details) (Aboalela *et al*., 2015; Alhareeri *et al*., 2020). However, there are limited methods to assess how the correlations between these exposures are associated with cancer treatment and how they evolve throughout the treatment and recovery process. Traditional analyses provide cross-sectional correlation estimates at individual time points, without taking the longitudinal structure into account (Prince *et al*., 2023). Despite advances in methods for longitudinal data, popular regression models such as generalized estimating equations (GEE) and generalized linear mixed-effects models (GLMM) cannot be directly applied to such correlation analysis. In addition, extending these models to Spearman’s rho is particularly challenging because these methods focus on modeling moments of variables and do not apply to Spearman’s rho defined by rankings of observations. As longitudinal studies have become the norm in modern clinical and laboratory research, there is a growing need for methods that can apply Spearman’s rho to such study data to facilitate analysis.

Some recent studies have extended Spearman’s rho to clustered data (Shih and Fay, 2017; Hunsberger *et al*., 2022; Tu, Li and Shepherd, 2025). Although longitudinal data is a type of clustered data, these methods only produce a single total correlation estimate pooled across time, and do not capture the temporal changes of the correlation throughout the study. Moreover, they are not developed based on a regression framework, thus not allowing for covariate adjustments. Liu et al. proposed a probability-scale residuals method, analogous to the partial Pearson’s correlation, to enable covariates adjustment, but this method is only applicable to cross-sectional data (Liu *et al*., 2018; Baba, Shibata and Sibuya, 2004). While Rosner and Glynn proposed a regression-based method for Spearman’s rho in clustered data, they used the relationship between the Pearson correlation and Spearman’s rho under bivariate normal distribution, which is often violated in practice (Rosner and Glynn, 2017). More importantly, due to the nature of longitudinal data collection, these datasets often contain missing values. Yet none of these methods consider how to account for the missing information, which may lead to biased results, if ignored. Although D’Angelo et al. proposed an EM algorithm based approach to handle missing data, their method is limited to crosssectional data and Pearson correlation (D’Angelo, Luo and Xiong, 2012).

To address these gaps and facilitate collaborative studies, we propose a novel framework to study the longitudinal association between metabolite profiles and mental wellness in BC patients receiving chemotherapy. We leverage a class of semiparametric models for between-subject attributes (Chen *et al*., 2016; Liu et al., 2022, 2024), the functional response models (FRM), to characterize time-varying changes in Spearman’s rho between metabolites and mental health outcomes, while controlling for key demographic and lifestyle covariates. The proposed approach accommodates missing data under the missing at random (MAR) assumption. Since MAR is popular in clinical research, this approach not only fills a critical methodological gap but also addresses a prevalent issue in practice. For the EPIGEN study, this framework offers a robust approach to study the dynamic association between metabolite measures and psychological wellness after BC treatment, as well as across different subpopulations to inform personalized medicine. To the best of our knowledge, this is the first integrative regression framework to model longitudinal Spearman’s rho, capture the temporal change of this correlation measure, and allow for covariate adjustments under MAR.

The paper is organized as follows. Section 2 introduces the study that motivates the method development. Section 3 describes the FRM and its extension to model Spearman’s rho for longitudinal data. Section 4 provides details on inference procedures for both complete and missing data. Section 5 conducts simulation studies to evaluate the proposed method. Section 6 applies the proposed approach and analytic framework to the motivating study. Section 7 gives concluding remarks and future recommendations.

## 2. Data Motivation

Our motivating data is from a one-year longitudinal study as part of the EPIGEN study. The study contains 77 women with early stage (I to IIIA) breast cancer from 5 regional cancer centers in Central Virginia (Aboalela *et al*., 2015; Alhareeri et al., 2020). The primary aim was to elucidate the complex relationships among psychoneurological symptoms (PNS; including anxiety, depression, fatigue, cognitive dysfunction, sleep disturbance, and pain), cancer treatment, the disease itself, and the combined influence of biological and psychological factors. Patients were followed for four time points in this longitudinal study: prior to chemotherapy (T1), at the time of their 4th cycle of chemotherapy treatment (T2), six months after the initiation of chemotherapy (T3), and one year after the initiation of chemotherapy (T4). Blood specimens were collected at each visit from each study participant and transported to the laboratory and stored using standard laboratory protocols. Blood samples were processed at the University of Florida’s Southeast Center for Integrated Metabolomics. Metabolomic profiling was performed using a Thermo Q-Exactive mass spectrometer coupled with a Dionex 3000 UHPLC system, operated in both positive and negative electrospray ionization modes, with additional details described in Lyon et al. (2023). PNS were evaluated at each time point over all study periods. For demonstration purposes, we mainly focus on the anxiety symptom, which was assessed using the Hospital Anxiety and Depression Scale (HADS) (Starkweather *et al*., 2017). The HADS is a 14-item self reported measure (each item scored 0-3) designed to assess anxiety and depressive symptoms, with demonstrated reliability and validity in women with BC (Snaith, 2003). The anxiety subscale ranges from 0 to 21, with scores of 6 or higher traditionally indicating clinically relevant symptoms in cancer patients (Singer *et al*., 2009). Demographic and lifestyle covariates, including race, BMI, current alcohol use, and current smoking status, were also collected. Our goal is to model the longitudinal rank correlation of metabolite levels and anxiety in BC patients receiving chemotherapy, and to examine racial disparities and temporal changes on the correlations while adjusting for these covariates. The metabolites identified through this analysis will be used in subsequent pathway analyses, such as metabolite set enrichment analysis, and for potential validation in independent cohorts to inform biomarker development and targeted interventions (Xia and Wishart, 2010; Lyon et al., 2023).

## 3. Functional Response Models for Spearman’s rho

In this section, we first provide a brief review of Spearman’s rho and in particular discuss some of its properties that are critical for formulating this correlation measure using the functional response model (FRM). We then focus on the extension of this correlation measure to longitudinal data under the FRM framework.

### 3.1 Spearman’s rho

Consider a cross-sectional study with *n* subjects and let **z**_*i*_ = (*u*_*i*_, *v*_*i*_)^⊤^ denote a bivariate continuous outcome (1 ⩽ *i* ⩽ *n*). Let *p*_*i*_ (*q*_*i*_) denote the rankings of *u*_*i*_ (*v*_*i*_) (1 ⩽ *i* ⩽ *n*), and 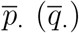 denote the mean rank scores of *p*_*i*_ (*q*_*i*_). Spearman’ rho, 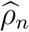, is defined by (Spearman, 1904):

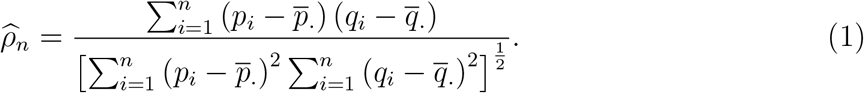

It is clear that 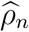 is the Pearson correlation applied to the rankings (*p*_*i*_, *q*_*i*_) of, rather than, the original variables **z**_*i*_, and thus can only be defined within a given study sample.

Like the Pearson correlation, 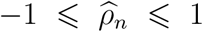, with 1 (−1) indicating perfect positive (negative) rank correlation. However, unlike the Pearson correlation, no linear relationship is required for *u*_*i*_ and *v*_*i*_, and 1 (−1) reflects perfect concordance (discordance) among pairs of **z**_*i*_ and **z**_*j*_ in the following sense:

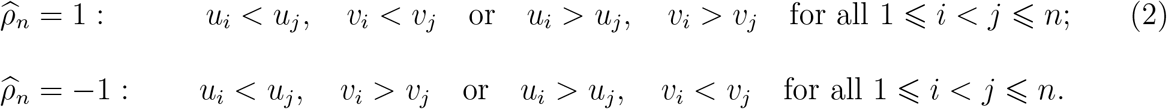

Thus, 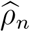 provides a sensible measure of correlation that captures non-linear monotonic association between *u*_*i*_ and *v*_*i*_.

Spearman’s correlation can also be expressed as a function of the estimated concordance for randomly drawn triplets of observations **z**_*i*_, **z**_*j*_ and **z**_*k*_. We define a triplet kernel

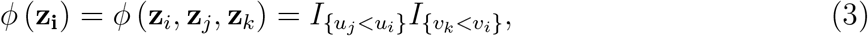

where *I*_{*x<y*}_ denotes an indicator that equals 1 if *x < y* and 0 otherwise. Then the population parameter for Spearman’s rho, denoted *ρ*, can be defined using this kernel function as *ρ* = 12*E* [*ϕ* (**z**_*i*_, **z**_*j*_, **z**_*k*_)] − 3, which is equivalent to the grade correlation in the case where *u* and *v* are continuous. (Kruskal, 1958). This formulation is important for our extension to longitudinal data. Theorem 1 shows that 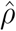 defined in (1) is a consistent estimator of such *ρ* and describes some basic properties of *ρ* as a correlation function. A more general version that addresses ties is included in the Supplementary materials along with the proof.

#### Theorem 1

*Let ϕ* (**z**_**i**_) *be defined as in* (3). *Then, we have:*

a. 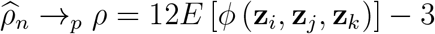
b. −1 ⩽ *ρ* ⩽ 1;
c. *ρ* = 1 (−1) *if and only if* **z**_*i*_ *has perfect-concordance (discordance), i.e*., *any pair of* **z**_*i*_ *and* **z**_*j*_ *are concordant (discordant), as defined in* (2);
d. *ρ* = 0 *if u and v are independent*.

### 3.2 Functional Response Models

Existing semiparametric regression models are defined based on a single-subject response. For example, let *y*_*i*_ and **x**_*i*_ denote some response and a vector of explanatory variables (predictors, covariates, independent variables) (1 ⩽ *i* ⩽ *n*). The most popular generalized linear model (GLM) is defined by 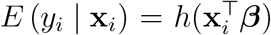, where *E* (*y*_*i*_ | **x**_*i*_) denotes the conditional mean of *y*_*i*_ given **x**_*i*_, *h*(·) is some specified function mapping 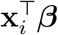 to the mean scale, and ***β*** is a vector of parameters. In this model, the response is a single-subject variable *y*_*i*_.

One inherent weakness of such single-subject-response-based regression models is their limited applications to modeling within-subject attributes. In modern biomedical applications, many outcomes of interest often involve between-subject relationships such as the mean of *ϕ* (**z**_*i*_, **z**_*j*_, **z**_*k*_) defined in Theorem 1. Such between-subject attributes cannot be readily accommodated within the traditional within-subject regression paradigm.

Therefore, we apply FRM to address these limitations (Chen *et al*., 2016; Liu et al., 2022, 2024). FRM is a class of semiparametric regression frameworks that model the conditional mean of a functional response involving outcomes of multiple subjects. Unlike the semiparametric GLMs or restricted moment models (Tsiatis, 2006), the conditional mean function *h* (·, ·) in an FRM is defined on explanatory variables from multiple subjects. Consequently, *h* (·, ·) describes how the conditional expectation of the functional response varies as the explanatory variables change. Specifically, FRM assumes

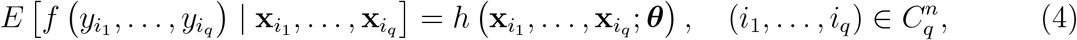

where *f* (·) is some function to capture the between-subject relationship of outcomes, *h* (·) is some smooth function (i.e., continuous second-order differentiable), 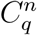 denotes the set of 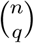 combinations of *q* distinct elements (*i*_1_, …, *i*_*q*_) from the integer set {1, …, *n*} and ***θ*** is a vector of parameters. By generalizing the response variable in this fashion, FRM is uniquely positioned to model *ϕ* (**z**_*i*_, **z**_*j*_, **z**_*k*_) defined by three individual-level responses **z**_*i*_, **z**_*j*_ and **z**_*k*_.

### 3.3 Functional Response Models for Spearman’s ρ

#### 3.3.1 Cross-sectional Data

Let {(**z**_*i*_, **x**_*i*_); 1 ⩽ *i* ⩽ *n*} be the observed cross-sectional data. Denote 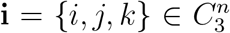, **z**_**i**_ = {**z**_*i*_, **z**_*j*_, **z**_*k*_}, and **x**_**i**_ = {**x**_*i*_, **x**_*j*_, **x**_*k*_}. Consider the functional response *ϕ* (**z**_**i**_) in (3). We are interested in how the Spearman’s rho between *u* and *v* varies with explanatory variables **x**_**i**_. Specifically, we introduce a covariate-specific parameter *ρ*(**x**) ∈ (−1, 1), and write *ρ*_**i**_ := *ρ*(**x**_**i**_) for the triplet **i**. We connect *ρ*_**i**_ to the triplet kernel by setting 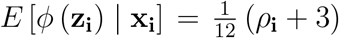, so we model *ρ*_**i**_ as a regression function of **x**_**i**_ through the link function 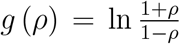 to map *ρ*_**i**_ to the real line. Thus we can define an FRM to model regression relationships of *ρ*_**i**_ with **x**_**i**_:

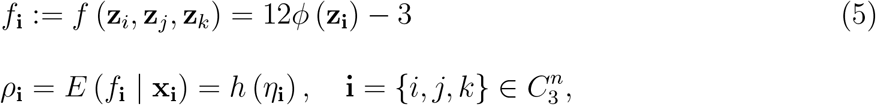

where 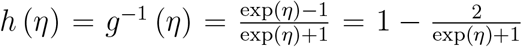 and *η*_**i**_ = *η*(**x**_**i**_, ***β***) ∈ (−∞, ∞) is a function of **x**_**i**_ parametrized by ***β***. For most applications, we use a linear *η*_**i**_ such that *η*_**i**_ = ***β***^⊤^***ξ***(**x**_**i**_) with ***ξ***(**x**_**i**_) being a function of **x**_*i*_, **x**_*j*_, **x**_*k*_. Since *h* (*η*) is a monotone increasing function of *η*, we interpret ***β*** according to whether ***β***^⊤^***ξ***(**x**_**i**_) increases or decreases as **x**_**i**_ changes.

For example, suppose we are interested in the association between a continuous variable, age, and the correlation measure. We can define a covariate age as the average or median of three individual ages, 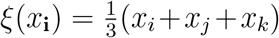, or *ξ*(*x*_**i**_) = median of *x*_*i*_, *x*_*j*_, *x*_*k*_, with *x*_*i*_ denoting the age of the *i*th subject. For a categorical covariate, consider comparing Spearman’s *ρ* between two groups defined by a binary indicator *x*_*i*_ ∈ {0, 1}. We can define ***ξ***(*x*_**i**_) as a vector of three binary indicators:

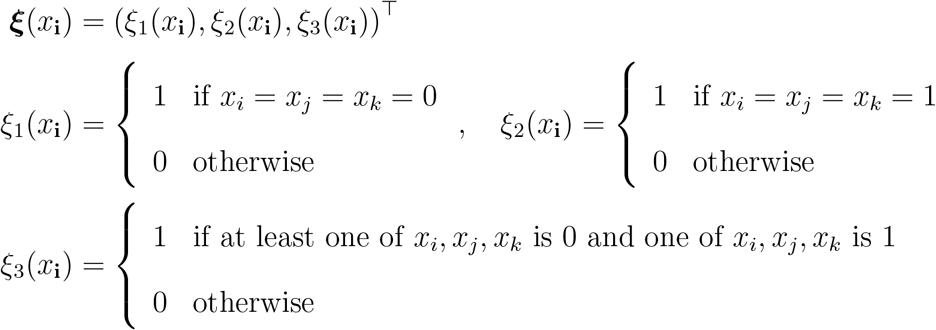

Then *η*_**i**_ = ***β***^⊤^***ξ***(*x*_**i**_) = *β*_0_*ξ*_1_(*x*_**i**_) + *β*_1_*ξ*_2_(*x*_**i**_) + *β*_2_*ξ*_3_(*x*_**i**_) and the Spearman’s rho is given by

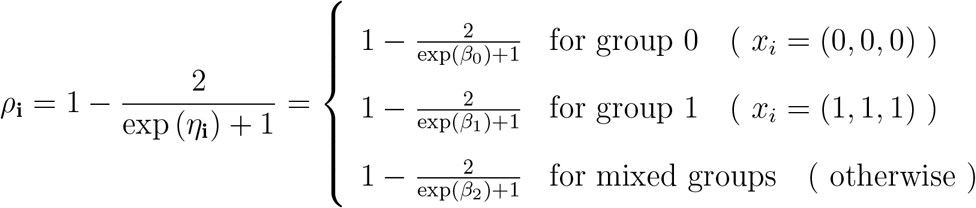

We can test the null of equal *ρ* between the groups by: *H*_0_ : *β*_1_ = *β*_0_. In this example, we are not interested in interpreting *β*_2_ as it pertains to the correlation defined by subjects’ outcomes from both groups. However, we may also define an alternative FRM that excludes triplets containing subjects from both groups (see Supplementary Materials for details).

#### 3.3.2 Longitudinal Data

By framing Spearman’s rho under FRM, we can readily extend the model in (5) to longitudinal data. Consider a longitudinal study with *m* assessments, indexed by *t* (1 ⩽ *t* ⩽ *m*). By applying the FRM in (5) for each *t*, we obtain:

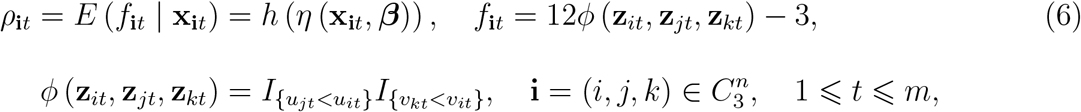

where **x**_**i***t*_ is a vector of time-varying explanatory variables including visit indicators and ***β*** is a vector of parameters and *η*_**i***t*_ := *η*(**x**_**i***t*_, ***β***) ∈ (−∞, ∞) is a function of **x**_**i**_ parametrized by ***β*** (1 ⩽ *t* ⩽ *m*). As in cross-sectional data case, we use a linear predictor *η*_**i***t*_ = ***β***^⊤^***ξ***(**x**_**i***t*_) with ***ξ***(**x**_**i***t*_) being a function of **x**_*it*_, **x**_*jt*_, **x**_*kt*_ for most applications.

Using (6), we can readily test hypotheses concerning temporal trends in *ρ* over time through specifying *η* (**x**_**i***t*_, ***β***). For example, to model changes in *ρ*_*t*_ over time, we can define ***ξ***(**x**_**i***t*_) as a vector of indicators: ***ξ***(**x**_**i***t*_) = (*ξ*_2_, *ξ*_3_, · · ·, *ξ*_*m*_)^⊤^ = (*I*_{*t*=2}_, *I*_{*t*=3}_, · · ·, *I*_{*t*=*m*}_)^⊤^ with *I*_{*t*=*s*}_ an indicator function of *t* = *s*. Then the linear predictor has the form:

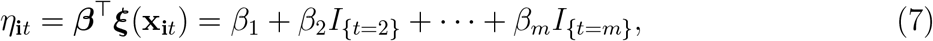

and the Spearman’s rho at each assessment time *t* is 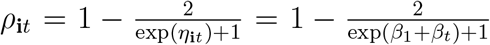. We can test the null of no temporal change in *ρ* by: *H*_0_ : *β*_*t*_ = 0 for all 2 ⩽ *t* ⩽ *m*. We can also add additional covariates *c*_**i**_ using additional linear terms *ξ*(*c*_**i**_) to (7).

## 4. Inference

Since cross-sectional data is a special case of longitudinal data, we focus on the latter for space consideration. We start with inference for complete data.

### 4.1 Inference Under Complete Data

Let

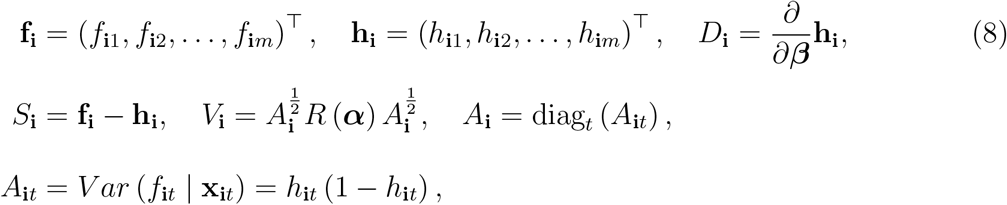

where diag_*t*_ (*A*_**i***t*_) denotes a diagonal matrix with *A*_**i***t*_ on the *t*-th diagonal and *R* (***α***) is a working correlation matrix parameterized by ***α*** to account for correlations among the repeated *f*_**i***t*_, akin to working correlations in GEE for clustered data (Liang and Zeger, 1986; Tang, He and Tu, 2023). We estimate ***β*** through the following U-statistics based Generalized Estimating Equations (UGEE):

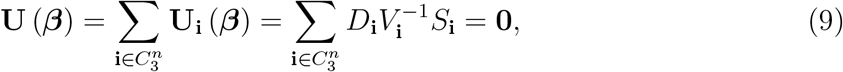

where *S*_**i**_ = (**f**_**i**_ − **h**_**i**_ (***β***)). Except for *R* (***φ***), the UGEE above is well-defined. If ***φ*** is unknown as in most applications, it must be estimated before solving the UGEE in (8). Although the consistency of 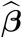 is independent of how ***α*** is estimated, the asymptotic normality of 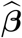 is guaranteed only when 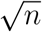-consistent estimators of ***α*** are used (Kowalski and Tu, 2008). The theorem below summarizes the asymptotic properties of the estimator from (9) (see Supplementary Materials for a proof).

#### Theorem 2

*Let*

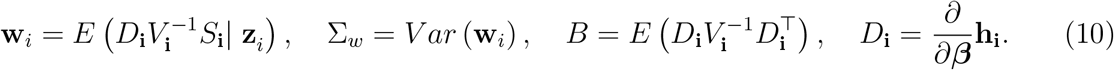

*Denote the UGEE estimate from solving* (9) *as* ***β***. *Then, under mild regularity conditions*,

a. 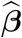 *is consistent;*
b. *if* 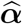 *is* 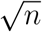*-consistent*, 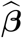 *is asymptotically normal, i.e*.,

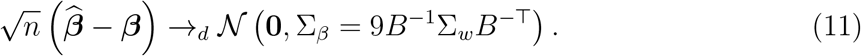

We can use this asymptotic normal distribution for inference about ***β***. Although Σ_*β*_ may be difficult to evaluate in closed form, it is readily estimated by replacing *B* and Σ_*w*_ each with a consistent estimator. A consistent estimator of *B* is given by 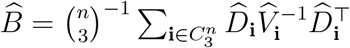, where 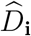 and 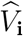 denotes *D*_**i**_ and *V*_**i**_ evaluated at 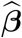. To estimate Σ_*w*_, we can first estimate **w**_*i*_ by fixing index *i* and enumerate combinations of the pair (*j, k*): 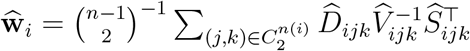, where 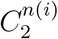 denotes the set of 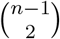 combinations of 2 distinct elements (*j, k*) from the integer set {1, …, *i* − 1, *i* + 1, …, *n*}. A consistent estimator of Σ_*w*_ and thus Σ_*β*_ are then given by:

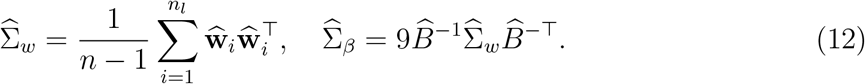

### 4.2 Inference under Missing Data

In the presence of missing data, the UGEE in (9) generally yields biased estimates, unless missing data follows the missing completely at random (MCAR) mechanism. To address the more general missing at random mechanism (MAR) mechanism, define a vector of missing (or rather observed) value indicators as follows:

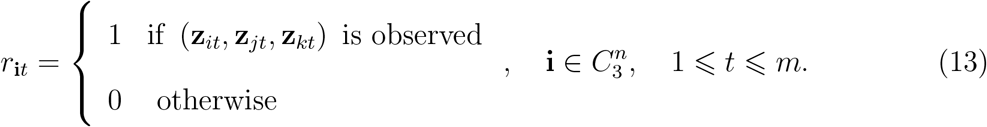

We assume no missing data at baseline *t* = 1 such that *r*_**i**1_ ≡ 1 for all 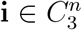.

Let 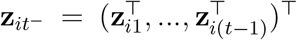, where 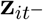 contains all **z**_*is*_ prior to time *t* (1 ⩽ *s* ⩽ *t* − 1). Under the monotone missing data pattern (MMDP) (Kowalski and Tu, 2008; Little and Rubin, 2019), 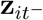 is observed, if **z**_*i*(*t*−1)_ is observed. Under MAR, we have 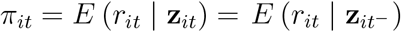, 2 ⩽ *t* ⩽ *m*. Thus, the last expression above is well-defined under MMDP.

Now let 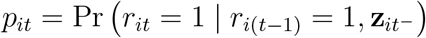, the one-step transition probability for observing the response from time *t* − 1 to *t*. We model *p*_*it*_ using logistic regression:

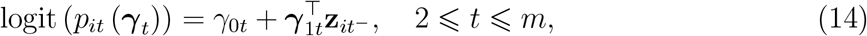

Where 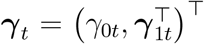. By testing ***γ***_1*t*_ = 0, we can evaluate whether missingness at time *t* is independent of prior outcomes. If ***γ***_1*t*_ ≠ 0, then the missingness depend on past outcomes, indicating the MAR mechanism. Under MMDP, for 2 ⩽ *t* ⩽ *m* and 1 ⩽ *i* ⩽ *n*, we have *π*_*it*_ (***γ***) = Pr (*r*_*it*_ = 1, *r*_*i*(*t*−1)_ = 1 | **z**_*it*_) = *p*_*it*_ 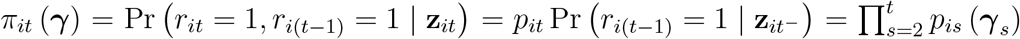, where 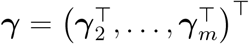, which provides a formula to estimate *π*_*it*_ using *p*_*it*_ modeled from (14).

Integrating the missing data model into (6) yields a new FRM as follows:

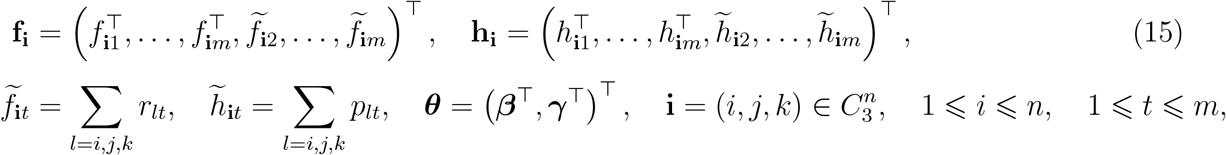

where **f**_**i***t*_ and **h**_**i***t*_ are defined in (6), *p*_*it*_ is given in (14) and *r*_*it*_ = 1 if **z**_*it*_ is observed. For notation brevity, denote ∑_*l*=*i,j,k*_ *a*_*l*_ = *a*_*i*_ + *a*_*j*_ + *a*_*k*_. For inference, let

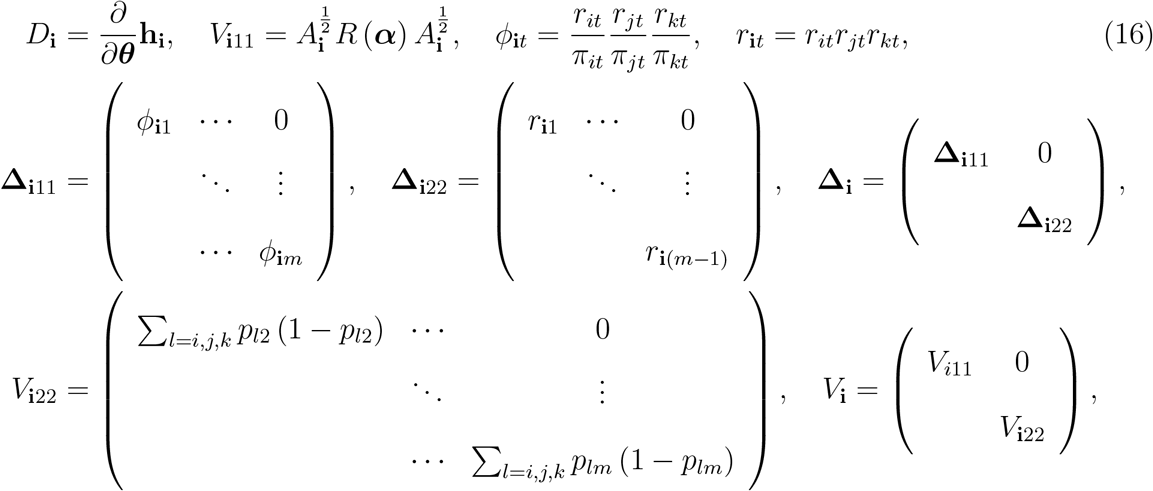

where *A*_**i**_ is defined in (8). We estimate ***θ*** from the following U-statistics-based weighted generalized estimating equations (UWGEE):

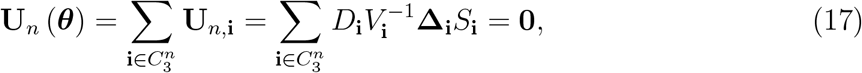

where *S*_**i**_ = (**f**_**i**_ − **h**_**i**_ (***β***)). The theorem below summarizes the asymptotic properties of the estimator of (17) (see Supplementary Materials for a proof).

#### Theorem 3

*Let* 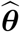 *denote the UWGEE estimate from solving the estimating equations in* (17) *upon substituting some consistent estimate* 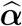. *Let*

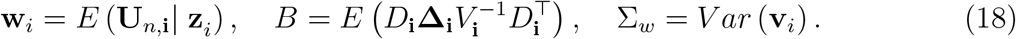

*Then, under under mild regularity conditions*, 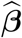 *is consistent and asymptotically normal*, 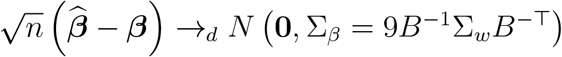.

As in Theorem 1, a consistent estimate of Σ_*β*_ is given by:

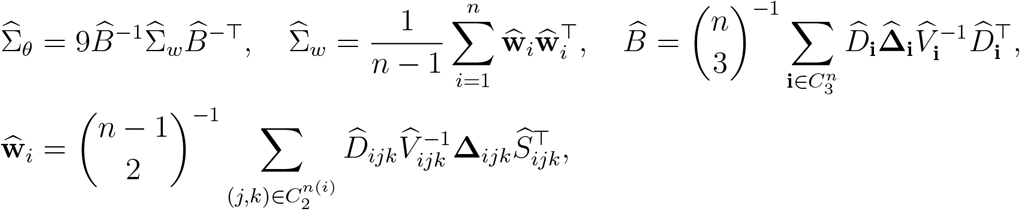

where 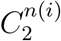 denotes the set of 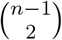 combinations of 2 distinct elements (*j, k*) from the integer set {1, …, *i* − 1, *i* + 1, …, *n*}.

## 5. Simulation Study

We use Monte Carlo simulation to examine the performance of the proposed approach by setting *n* = 50 (small), 150 (medium) and 500 (large) sample sizes. We set the Type-I error *α* = 0.05 and Monte Carlo sample size *M* = 1, 000 in all analyses unless stated otherwise.

### 5.1 Complete Data

We start with complete longitudinal data to evaluate the performance of our proposed FRM-based-approach for modeling temporal changes in Spearman correlations over time. The underlying data generating mechanism assumes that each subject *i* has bivariate measurements (*u*_*it*_, *v*_*it*_) at times *t* = 1, 2, 3. Define **z**_*i*_ = (*u*_*i*1_, *v*_*i*1_, *u*_*i*2_, *v*_*i*2_, *u*_*i*3_, *v*_*i*3_)^⊤^ and let **z**_*i*_ ∼ *N* (*µ*, Σ) with *µ* = (1, 1, 1, 1, 1, 1)^⊤^ and the structured variance matrix Σ with unit variances and include both within-time dependence (*δ* = 0.5, representing the correlation between *u*_*it*_ and *v*_*it*_) and between-time dependence (*ϕ* = 0.25, representing the correlation between *u*_*is*_ and *u*_*it*_). The correlation between *u*_*is*_ and *v*_*it*_ is also set to be *ϕ* = 0.25.

In this setting, we have two binary indicators for the 2nd and 3rd assessments: *x*_**i***t*,2_ = *I*_{*t*=2}_ and *x*_**i***t*,3_ = *I*_{*t*=3}_. Thus the linear predictor is *η*_**i***t*_ = *β*_1_ + *β*_2_*x*_**i***t*,2_ + *β*_3_*x*_**i***t*,3_ = *β*_1_ + *β*_2_*I*_{*t*=2}_ + *β*_3_*I*_{*t*=3}_. The Spearman’s rho at each time is

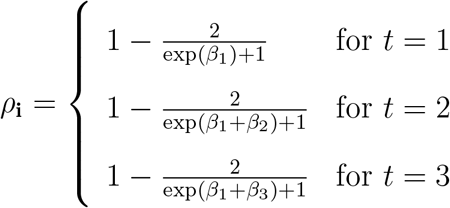

In this simulation, we set *β*_2_ = *β*_3_ = 0 (additional simulations with different *β*’s that leads to time-varying Spearman’s rho are included in Supplementary Materials). Given the formula derived in Heinen and Valdesogo (2020), the true value for Spearman’s rho is 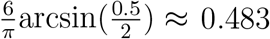, while the true value for 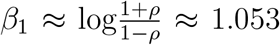. We apply the FRM model where *β*_1_ represents the baseline log-odds correlation, while *β*_2_ and *β*_3_ capture temporal changes from baseline to time point *t* = 2 and *t* = 3, respectively. We assess asymptotic performance by comparing asymptotic and empirical variance estimates for *β*_1_, *β*_2_, and *β*_3_. We also evaluate the performance of the approach relevant to three null hypotheses: (1) *β*_2_ = 0, (2) *β*_3_ = 0, (3) *β*_2_ = *β*_3_ = 0.

Table 1(a) shows the estimates for ***β***, asymptotic and empirical standard error, and the estimates for Spearman’s rho (*ρ*). The baseline parameter *β*_1_ was consistently estimated across all sample sizes, with empirical standard errors close to their asymptotic counterparts. The estimated Spearman correlations 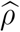 were close to the true value (≈ 0.483) across all time points. Estimates of *β*_2_ and *β*_3_ were close to the true values (0), and asymptotic and empirical standard errors were also close. The corresponding Type-I error rates shown in Table 1(b) were close to the nominal level of 0.05, even for small sample size case (*n* = 50).

**Table 1:** Simulation Results for Longitudinal Complete Data.

(a) Estimates of $\beta$ and $\rho$
| Time point | $\hat{\beta}$ | Asymptotic $SE(\hat{\beta})$ | Empirical $SE(\hat{\beta})$ | $\hat{\rho}$ |
| --- | --- | --- | --- | --- |
| $n = 50$ | | | | |
| $t = 1 (\beta_1)$ | 1.049 | 0.303 | 0.313 | 0.481 |
| $t = 2 (\beta_2)$ | 0.017 | 0.419 | 0.419 | 0.488 |
| $t = 3 (\beta_3)$ | 0.032 | 0.419 | 0.424 | 0.493 |
| $n = 150$ | | | | |
| $t = 1 (\beta_1)$ | 1.061 | 0.171 | 0.171 | 0.486 |
| $t = 2 (\beta_2)$ | -0.001 | 0.236 | 0.239 | 0.485 |
| $t = 3 (\beta_3)$ | 0.006 | 0.236 | 0.239 | 0.488 |
| $n = 500$ | | | | |
| $t = 1 (\beta_1)$ | 1.056 | 0.093 | 0.093 | 0.484 |
| $t = 2 (\beta_2)$ | -0.004 | 0.128 | 0.128 | 0.482 |
| $t = 3 (\beta_3)$ | 0.001 | 0.128 | 0.126 | 0.484 |

(b) Type-I Error Rates
| Sample Size | $H_0 : \beta_2 = 0$ | $H_0 : \beta_3 = 0$ | $H_0 : \beta_2 = \beta_3 = 0$ |
| --- | --- | --- | --- |
| $n = 50$ | 0.052 | 0.058 | 0.063 |
| $n = 150$ | 0.045 | 0.050 | 0.044 |
| $n = 500$ | 0.054 | 0.046 | 0.050 |

### 5.2 Missing Data

Assume we still follow the data generating process as in Section 5.1. In addition, we implement MMDP as explained in Section 4.2. We specify the MAR mechanism where the probability of missing future observations depends on previously observed values. Specifically, at *t* = 1, all observations are complete (*r*_*i*1_ = 1, *i* = 1, …, *n*). For *t* = 2, 3, the missing indicator follow sequential logistic models:

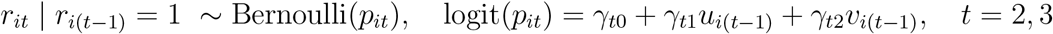

Let ***γ***_*t*_ = (*γ*_*t*0_, *γ*_*t*1_, *γ*_*t*2_)^⊤^ for *t* = 2, 3, and 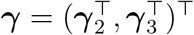. To satisfy MMDP, if *r*_*it*_ = 0, then *r*_*i*(*t*−1)_ = 0. The intercept parameters *γ*_20_ and *γ*_30_ are chosen to achieve target observation rates of 85% at *t* = 2 and 75% at *t* = 3. Let *π*_*i*1_ = 1, *π*_*i*2_(***γ***) = *p*_*i*2_(***γ***), and *π*_*i*3_(***γ***) = *p*_*i*2_(***γ***)*p*_*i*3_(***γ***) denote the cumulative observation probabilities through each time point. We set *γ*_21_ = *γ*_22_ = *γ*_31_ = *γ*_32_ = 1 and solve 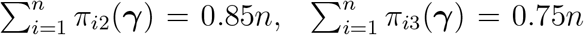 with *n* = 50, 000 to obtain *γ*_20_ = 0.541 and *γ*_30_ = 0.799.

We jointly estimate the correlation parameters ***β*** = (*β*_1_, *β*_2_, *β*_3_)^⊤^ and the missing data mechanism parameters ***γ*** using the UWGEE in (17). Table 2(a) shows the mean estimates of ***β***, the asymptotic and empirical standard errors, and the derived estimates for *ρ* under our monotone MAR missing pattern. The *β*_1_ and the estimated *ρ* values remained well-estimated, suggesting robustness of the UWGEE method to missingness. Estimates of *β*_2_ and *β*_3_ were still close to zero. Empirical and asymptotic standard errors remained close to each other. Type-I error rates (Table 2(b)) were also close to 0.05.

**Table 2:** Simulation Results for Longitudinal Data under MMDP.

(a): Estimates of $\beta$ and $\rho$
| Time point | $\hat{\beta}$ | Asymptotic $SE(\hat{\beta})$ | Empirical $SE(\hat{\beta})$ | $\hat{\rho}$ |
| --- | --- | --- | --- | --- |
| $n = 50$ | | | | |
| $t = 1$ | 1.058 | 0.304 | 0.301 | 0.484 |
| $t = 2$ | 0.024 | 0.445 | 0.453 | 0.493 |
| $t = 3$ | 0.000 | 0.470 | 0.481 | 0.485 |
| $n = 150$ | | | | |
| $t = 1$ | 1.067 | 0.171 | 0.174 | 0.488 |
| $t = 2$ | -0.006 | 0.256 | 0.258 | 0.486 |
| $t = 3$ | -0.031 | 0.263 | 0.269 | 0.476 |
| $n = 500$ | | | | |
| $t = 1$ | 1.056 | 0.093 | 0.095 | 0.484 |
| $t = 2$ | 0.005 | 0.140 | 0.143 | 0.486 |
| $t = 3$ | -0.020 | 0.143 | 0.146 | 0.476 |

(b): Type-I Error Rates
| Sample Size | $H_0 : \beta_2 = 0$ | $H_0 : \beta_3 = 0$ | $H_0 : \beta_2 = \beta_3 = 0$ |
| --- | --- | --- | --- |
| $n = 50$ | 0.062 | 0.060 | 0.056 |
| $n = 150$ | 0.049 | 0.065 | 0.058 |
| $n = 500$ | 0.052 | 0.061 | 0.054 |

## 6. Case Study

We applied our method to the EPIGEN longitudinal metabolomics dataset described in Section 2. At each visit, blood specimens were processed to quantify 2,395 metabolites. For each metabolite, our goal was to model its correlation with anxiety throughout chemotherapy and recovery, while adjusting for race, BMI, current alcohol use, and current smoking status. After excluding three participants with missing BMI, the final analytical sample size was 74. Across the 2,395 metabolites, most had low missing rates, with a median of 5.4% (IQR = 5.4% - 8.1%), although some metabolites had missing rates as high as 54.1%.

The study includes four visits in total, indexed by *t* = 1, …, 4. At each visit *t*, we construct the triplet-based response *f*_**i***t*_ from *ϕ*(**z**_*it*_, **z**_*jt*_, **z**_*kt*_) as in (6), where **z**_*it*_ = (*u*_*it*_, *v*_*it*_) denotes the paired outcomes of a metabolite measure and anxiety score for subject *i* at visit *t*. The FRM then specifies the conditional mean *E*(*f*_**i***t*_ | **x**_**i***t*_), where **x**_**i***t*_ includes visit indicators along with race and the lifestyle covariates. Since **x**_**i***t*_ is defined at the triplet level, we construct triplet-level covariates as follows: for the continuous covariate, BMI, we use the average of the three values (i.e., (BMI_*i*_ +BMI_*j*_ +BMI_*k*_)*/*3); for categorical covariates, we create dummy variables for each of the three elements and construct triplet-level indicators of categorical equality or dissimilarity across the three values, as described in Section 3.3.1. By including visit indicators in **x**_**i***t*_, the model allows metabolite-anxiety association to change over time. A special case of this framework that models time-constant correlation can be achieved by omitting the time indicator terms, but we allow for time variation as both metabolic measure and anxiety level may change during chemotherapy and recovery.

To address missing data, we examined the missing completely at random (MCAR) assumption for each metabolite. For metabolites where the MCAR assumption was not rejected, we applied the approach under MCAR; otherwise, we applied the approach under MAR. We model the time-varying correlations between each individual metabolite and the anxiety score, and examine whether the metabolite–anxiety associations change over time as individuals undergo chemotherapy and recovery, as well as whether the overall correlation differs by racial groups after adjustment under the additive model. Because of the large number of metabolites, we used a two-stage procedure to identify such metabolites. In the first step, we applied a filtering step based on coefficient p-values from the FRM, retaining metabolites with p-values *<* 0.2 for the covariate of interest. For temporal change from before and after the surgery, we screened on p-value for the visit 2 indicator, which retained 728 metabolites. For racial differences, we screened on p-values for the race indicator comparing White patients and Black/African American patients, which retained 742 metabolites. In the second step, we applied a Bonferroni correction within each screened set to adjust for multiple comparisons and determine which metabolites remained statistically significant.

### 6.1 Covariate-adjusted Spearman’s rho

Figure 1 presents the distribution of Spearman’s rho between metabolite profiles and Total Anxiety Score, both before and after covariate adjustments. Panel (a) shows the unadjusted correlations across all four timepoints, while panel (b) and (c) separately shows the magnitudes of Spearman’s rho between metabolites and anxiety for Black/African American patients and White patients, adjusting for smoking status, current alcohol use, and BMI. The top 20 metabolites for each race group (listed in Table 3), selected based on the mean absolute value of Spearman’s rho across all four time points, are highlighted in red. Generally, the magnitude of the estimated correlations are higher after controlling for covariates. This pattern suggests that if we do not control for demographic and lifestyle factors, the correlations between metabolites and anxiety may appear to be weaker. After adjustment, the estimated correlations become stronger and more scientifically meaningful.

**Table 3:**
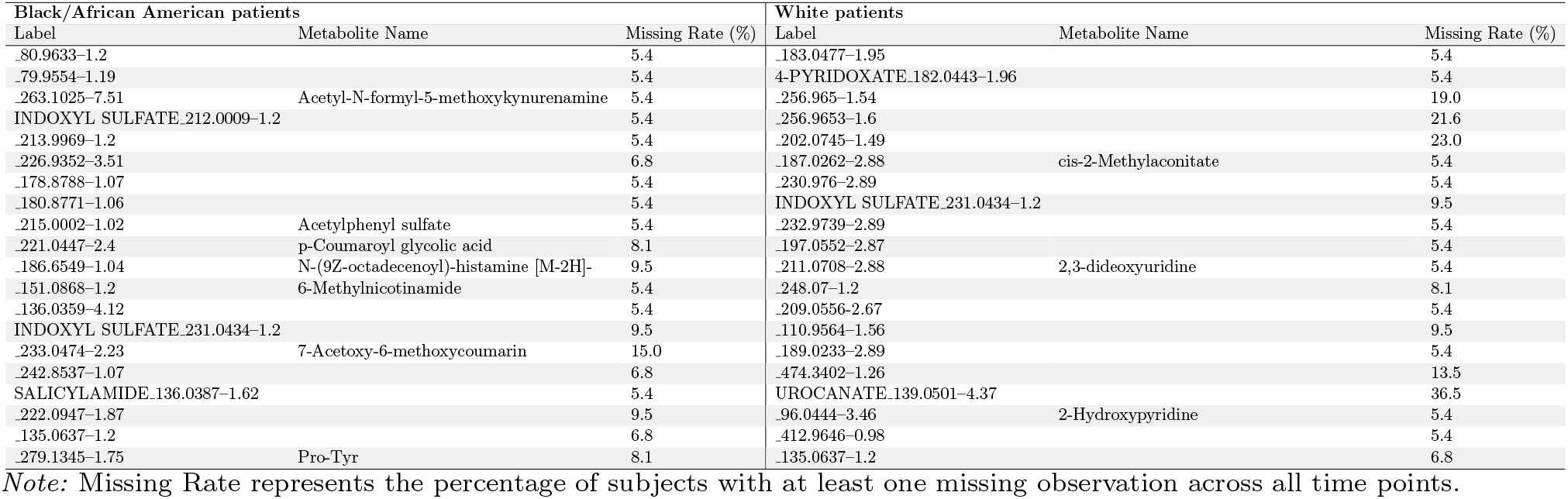
Top 20 metabolites with the largest magnitude of Spearman’s *ρ* with Total Anxiety Score.

**Figure 1:**
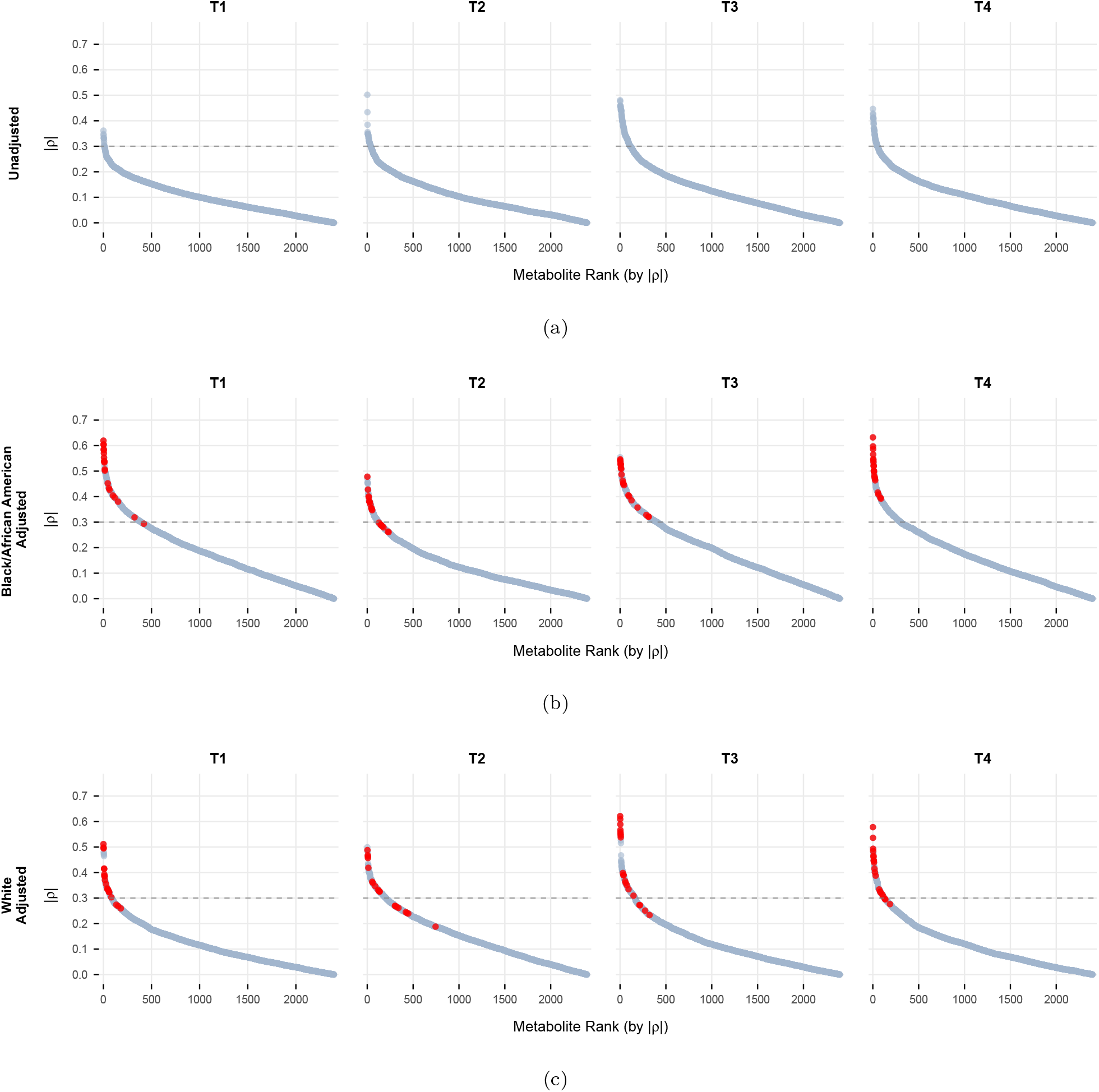
Absolute Value of Spearman’s *ρ* between all measured metabolites and Total Anxiety Score across four timepoints: (a) unadjusted estimates, (b) covariate-adjusted estimates for Black/African American patients, (c) covariate-adjusted estimates for White patients. The adjusted model includes race, smoking status, current alcohol use, and BMI. Top 20 metabolites with the strongest correlations with Total Anxiety Score are highlighted in red in the adjusted analyses.

Table 3 shows the set of metabolites most strongly correlated with anxiety for Black/African American patients and White patients respectively. The table includes the experimental labels and, when available, the corresponding metabolite names identified via The Human Metabolome Database (HMDB) and Metabolomics Workbench (Wishart *et al*., 2022; Metabolomics Workbench). While INDOXYL SULFATE 231.0434–1.2 and 135.0637–1.2 were among the top correlated metabolites in both groups, other metabolites showed strong correlations in one group but not the other. Our results suggests that the biological relationship between metabolites and anxiety may differ across racial groups, motivating further investigations on each group separately.

**Figure 2:**
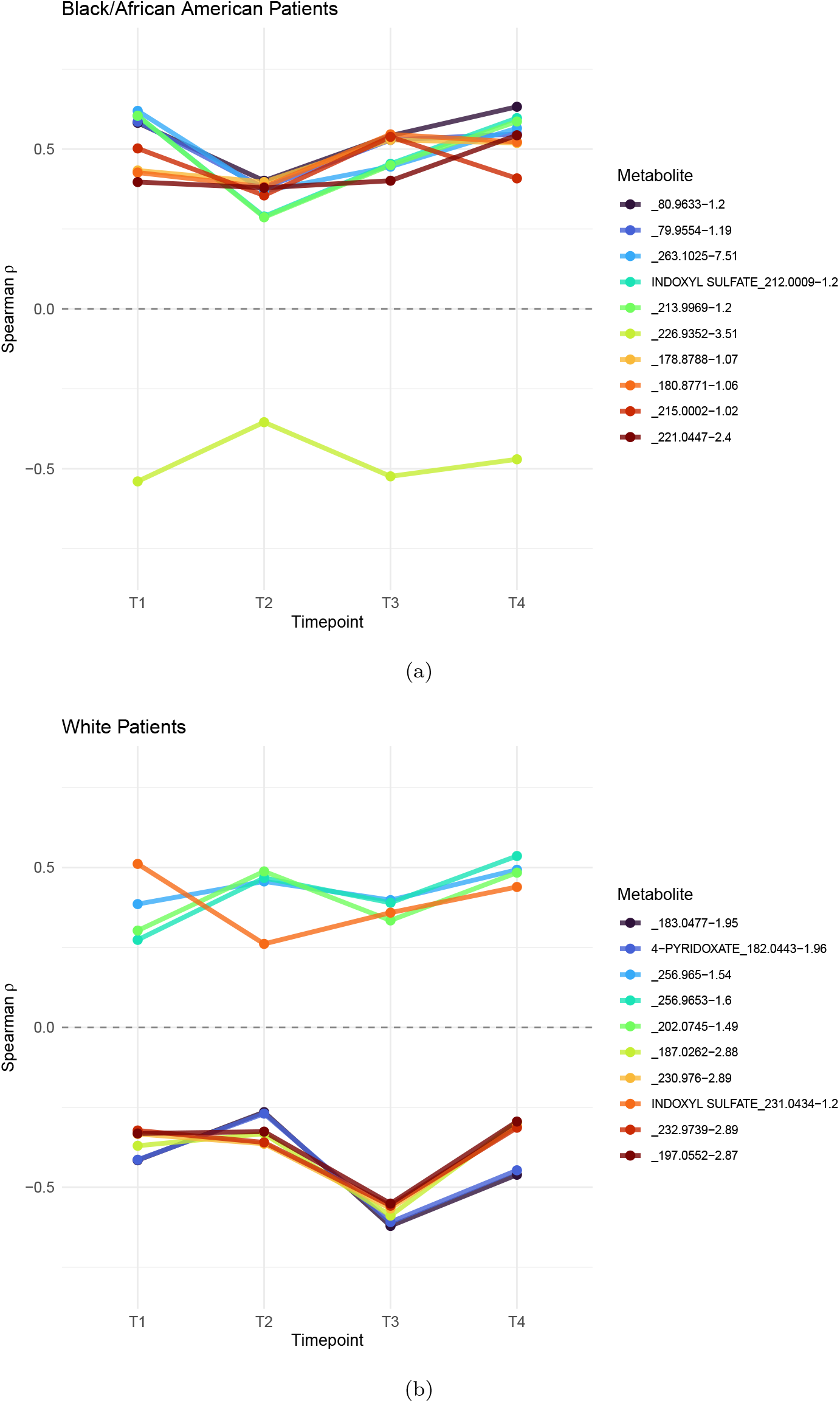
Top 10 metabolites associated with Total Anxiety Score based on the mean absolute value of Spearman’s *ρ* across all timepoints: (a) Top 10 for Black/African Patients (b) Top 10 for White Patients

### 6.2 Racial difference in Metabolite-Anxiety correlation

One big advantage of our FRM regression framework is that it allows hypothesis tests for regression coefficients. We tested the null hypothesis that the correlation between each metabolite and Total Anxiety Score was the same for Black/African American and White patients, adjusting for current smoking status, alcohol use, and BMI. In this application, we consider an additive model. After applying the two-step multiple-testing procedure for the race effect as described in Section 6, one metabolite, 826.5561–6.02 (identified as PE (40:4-2OH)), was found to have a significant racial difference in its correlation with anxiety (adjusted p = 0.0496). Figure 3(a) presents the estimated correlations for metabolite PE (40:4-2OH) with anxiety over the four time points. For this specific metabolite, the correlation was consistently positive among Black/African American patients, while it was consistently negative among White patients.

**Figure 3:**
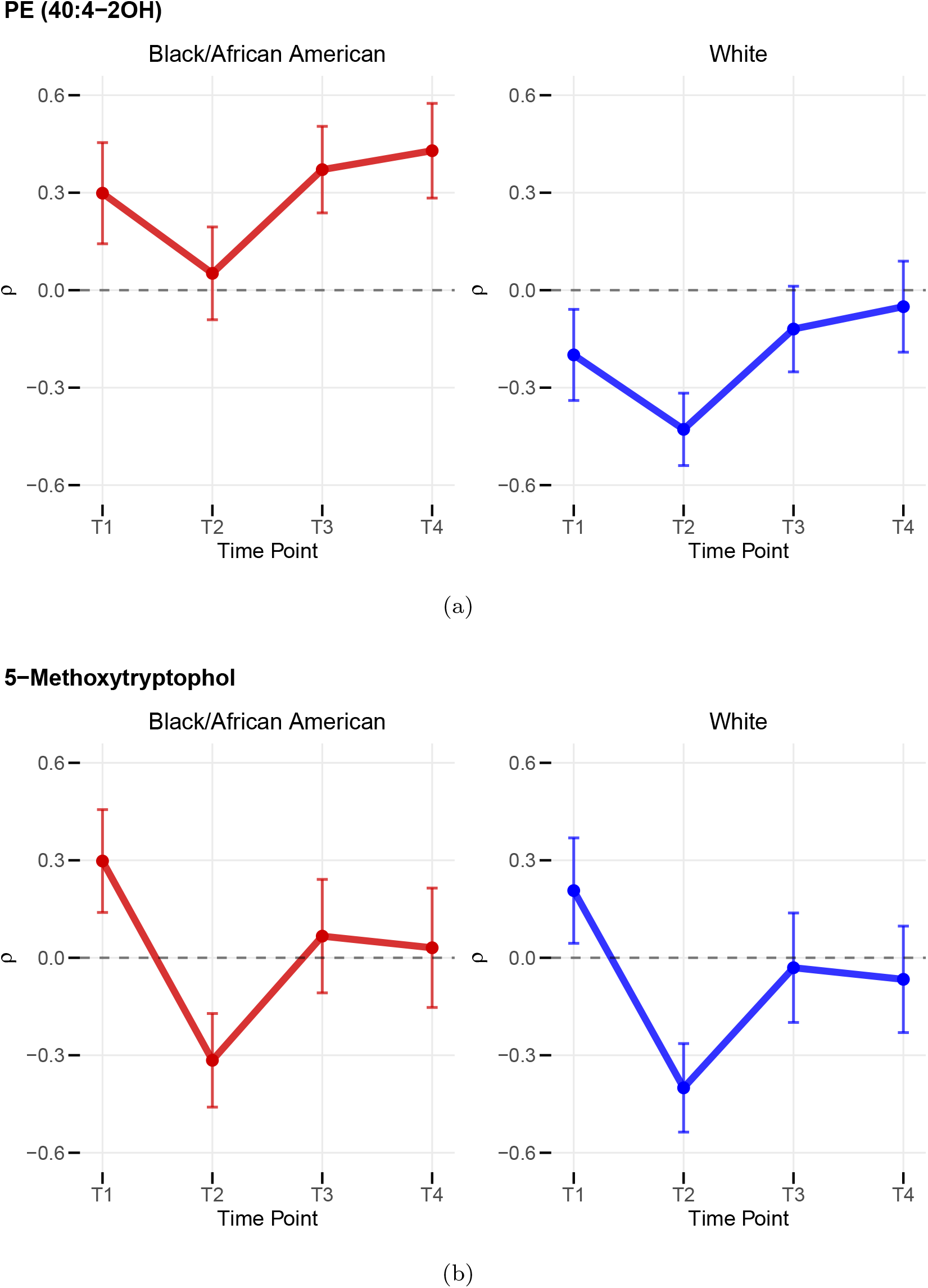
(a) Spearman’s *ρ* between PE (40:4-2OH) and Total Anxiety Score Spearman’s *ρ* between 5-Methoxytryptophol and Total Anxiety Score

### 6.3 Temporal changes in Metabolite-Anxiety correlation (T1 vs. T2)

In addition, we evaluated the time trend in the correlation between metabolite measures and Total Anxiety Score. Specifically, we tested the null hypothesis that the correlation between each metabolite and Total Anxiety Score was the same at baseline (T1, prior to chemotherapy) and at T2 (after the fourth chemotherapy treatment), using an additive model. After multiple testing adjustments for the visit 2 effect, one metabolite, 192.1021–1.72 (identified as 5-Methoxytryptophol), was identified as significant (adjusted p = 0.0351). Figure 3(b) shows the estimated correlations for metabolite 5-Methoxytryptophol with anxiety for both Black/African American and White patients. For this metabolite, the correlation decreased sharply after baseline for both racial groups. This pattern indicates that chemotherapy may affect the relationship between this metabolite and anxiety dramatically, from positive to negative, particularly during the treatment period. However, six months to one year after chemotherapy (T3, T4), such negative correlation weakens and may even revert to positive, although its magnitude remains smaller than at baseline (T1).

## 7. Discussion

In this paper, we developed and applied a regression framework to model the longitudinal Spearman’s rho between metabolites and anxiety levels among breast cancer patients who experienced chemotherapy treatment and recovery. By focusing on rank-based Spearman’s rho, the proposed method is inherently robust against non-linearity. With our proposed FRM framework, we are able to model changes in Spearman’s rho over time for longitudinal studies and allow for covariate-adjustments and missing data under MAR assumption. The method performs well, even when the sample size is as small as 50, as evidenced by the simulation study.

Although we focus on longitudinal data, the proposed approach extends naturally to other clustered settings. By replacing the repeatedly assessed outcomes **z**_*it*_ with appropriately defined variables, the FRM in (6) can also be used to model Spearman’s rho across clusters (Tu, Li and Shepherd, 2025). For example, if **z**_*i*1_ = (*u*_*i*_, *v*_*i*_)^⊤^ and 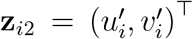 are two pairs of variables of interest from two clusters, the same modeling strategy allows us to test whether the correlation *ρ*_1_ between *u*_*i*_ and *v*_*i*_ differs from the correlation *ρ*_2_ between 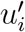 and 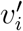. This demonstrates that our framework can be applied more broadly than the longitudinal setting.

When applied to the motivating EPIGEN study, our approach reveals that the correlation between anxiety level and various metabolites are generally stronger after controlling for race, current smoking status, current alcohol use, and BMI. We successfully identified sets of metabolites that are most strongly correlated with anxiety level separately for Black/African American patients and White patients, after controlling for important covariates. Additionally, our approach also enables hypothesis tests to assess the time-trends of metabolite–anxiety correlations and whether the correlations differ across covariate groups. Our analysis revealed significant racial differences in the correlation between PE (40:4-2OH) and anxiety level and a significant change in the correlation between 5-Methoxytryptophol and anxiety level after chemotherapy treatment.

The identified metabolites will be validated in independent cohorts in future studies. Metabolites that demonstrate strong and reproducible temporal associations with mental wellness may serve as diagnostic, prognostic, or monitoring biomarkers. They will also be incorporated into pathway analyses, such as metabolite set enrichment analysis, to identify biological pathways that are significantly up or down regulated, as described in (Lyon *et al*., 2023). Furthermore, these metabolites could inform targeted interventions, including pharmacological or nutritional strategies designed to modulate metabolite levels (Vernieri *et al*., 2016; Beger et al., 2016).

The proposed framework offers a powerful and innovative regression based approach for modeling longitudinal correlations using Spearman’s rho. It opens a new window for uncovering the dynamic relationships between biomarkers and disease before, during, and after treatment. In future studies, our team will apply this framework to further evaluate differential effects of chemotherapy on correlations and potential contributing factors, including dietary patterns, timing of chemotherapy, and other relevant variables.

Our current framework allows missing data to follow either MCAR or MAR mechanisms. In practice, such models might be mis-specified and lead to biased correlation estimations. To improve robustness against model mis-specification, we plan to extend the current work by developing doubly robust estimators (DRE) for longitudinal Spearman’s rho (Bang and Robins, 2005; Tsiatis, 2006).

When applied to ultra-high dimensional genomic or metabolomic data, the high-dimensionality can impose a substantial computation burden. In our analysis, we have 74 subjects, 2395 metabolites, and 4 time points, and the computation time was about 2 hours (to run all 2395 metabolites), facilitated by the use of RCPP. However, as the sample size and number of metabolites or genetic features increase, computation time may increase dramatically. Moreover, metabolites are often highly correlated within the same pathway, so running Spearman’s rho individually for each metabolite ignores these interdependencies and may lead to redundant results. Testing each metabolite separately also inflates the risk of false positives due to multiple comparisons. Given that our goal was to develop an integrative framework for modeling longitudinal Spearman’s rho, we did not implement advanced feature screening approaches in our real analysis (Fan and Lv, 2008; Li, Zhong and Zhu, 2012). Instead, we applied an ad hoc pre-selection approach based on p-value thresholding. Future work will continue to explore more robust feature screening strategies to improve computational efficiency and statistical interpretability in high-dimensional longitudinal data. Analysis code for the simulations and case study (although without the actual case study data) are available at https://github.com/yiwen99/Longitudinalspearman.

## Acknowledgments

Shepherd, B.E. is partially supported by NIH grant R01AI093234.

